# “Stretch-Growth” of Motor Axons in Custom Mechanobioreactors to Generate Long-Projecting Axonal and Axonal-Myocyte Constructs

**DOI:** 10.1101/598755

**Authors:** Kritika S. Katiyar, Laura A. Struzyna, Suradip Das, D. Kacy Cullen

**Affiliations:** Center for Brain Injury & Repair, Department of Neurosurgery, Perelman School of Medicine, University of Pennsylvania, Philadelphia, PA 19104; Center for Neurotrauma, Neurodegeneration & Restoration, Corporal Michael J. Crescenz Veterans Affairs Medical Center, Philadelphia, PA 19104; School of Biomedical Engineering, Science & Health Systems, Drexel University, Philadelphia, PA 19104; Department of Bioengineering, School of Engineering & Applied Science, University of Pennsylvania, Philadelphia, PA 19104

## Abstract

The central feature of peripheral motor axons is their remarkable lengths as they project from a motor neuron residing in the spinal cord to an often-distant target muscle. However, to date *in vitro* models have not replicated this central feature owing to challenges in generating motor axon tracts beyond a few millimeters in length. To address this, we have developed a novel combination of micro-tissue engineering and mechanically assisted growth techniques to create long-projecting centimeter-scale motor axon tracts. Here, primary motor neurons were isolated from the spinal cords of rats and induced to form engineered micro-spheres via forced aggregation in custom micro-wells. This three-dimensional micro-tissue yielded healthy motor neurons projecting dense, fasciculated axonal tracts. Within our custom-built mechanobioreactors, motor neuron culture conditions, neuronal/axonal architecture, and mechanical growth conditions were systematically optimized to generate parameters for robust and efficient “stretch-growth” of motor axons. We found that axons projecting from motor neuron aggregates were able to respond to axon displacement rates at least 10 times greater than that tolerated by axons projecting from dissociated motor neurons. The growth and structural characteristics of these stretch-grown motor axons were compared to benchmark stretch-grown axons from sensory dorsal root ganglion neurons, revealing similar axon densities yet increased motor axon fasciculation. Finally, motor axons were integrated with myocytes and then stretch-grown to create novel long-projecting axonal-myocyte constructs that better recreate characteristic dimensions of native nerve-muscle anatomy. This is the first demonstration of mechanical elongation of spinal cord motor axons and may have applications as anatomically inspired *in vitro* testbeds or as tissue engineered “living scaffolds” for targeted axon tract reconstruction following nervous system injury or disease.

**Significance Statement:** We have developed novel axon tracts of unprecedented lengths spanning either two discrete populations of neurons or a population of neurons and skeletal myocytes. This is the first demonstration of “stretch-grown” motor axons that recapitulate the structure of spinal motor neurons *in vivo* by projecting long axons from a pool of motor neurons to distant targets, and may have applications as anatomically inspired *in vitro* test beds to study mechanisms of axon growth, development, and neuromuscular function in anatomically accurate axo-myo constructs; as well as serve as “living scaffolds” *in vivo* for targeted axon tract reconstruction following nervous system trauma.

## Introduction

The peripheral nervous system (PNS) consists of nerves that project from the spinal cord to the periphery. These nerves are comprised of bundles of axons that stem from neuronal cell bodies housed adjacent to or within the spinal column and project to the rest of the body. For instance, sensory dorsal root ganglia (DRG) are located in the posterior (dorsal) root of the spinal cord and project axons to the periphery; they are responsible for sensory stimuli such as pain, temperature, and mechanical stimulus. Motor neurons are located in the ventral horn, within the gray matter of the spinal cord and project long axons through the ventral root that innervate distal muscles. In humans, motor neuron somata can be as large as 100 μm in diameter with axons projecting over 1 m to distal targets ^1^. During development, growing axons are guided to the appropriate end target by pathways formed by existing cells – glial “guidepost cells” as well as “pioneer” axons that have already reached the end target ^2^. After reaching end targets, these axon tracts are subjected to mechanical forces (e.g., tension or lengthening) as the body grows throughout development, resulting in a natural form of so-called “stretch-growth”. Indeed, based on the application these growth-promoting forces, peripheral axons can reach lengths that are thousands of times greater than the diameter of the neuronal cell body that sustains them.

However, *in vitro* models do not replicate this central feature of long-projecting axonal tracts owing to challenges in generating motor axon tracts beyond a few millimeters in length. This is likely due to current culture systems presenting suboptimal two-dimensional (2D) conditions that lack the necessary chemotactic, haptotactic, and mechanical interactions needed to support generation of long motor axon tracts. Therefore, many researchers are turning to 3D culture systems to more accurately mimic living systems ^3^-^5^. One form of 3D cell culture is creating cell “spheroids”. Spheroids are aggregates of cells that offer a high throughput way of modeling the complex morphology and physiology of *in vivo* tissue by allowing co-culture of various cell types on or within biomaterials to more accurately study cell-cell and cell-matrix interactions. However, the process of spheroid formation, also referred to as “self aggregation” or “forced aggregation”, has yet to be applied in conjunction with techniques to grow long-projecting (e.g., centimeter scale) axon tracts.

The axon growth process we employ is inspired by the phenomenon of axon stretch growth seen in development, allowing us to generate long axon tracts in custom-built mechanobioreactors. Indeed, this builds on the work of Smith and colleagues, who have demonstrated the controlled application of mechanical forces to produce stretch grown axons from a number of neuronal sources, including iPSC derived DRG, human cadaveric DRG, and DRG from embryonic rats, spanning several centimeters ^6-10^. Remarkably, this work demonstrated so-called “stretch-growth” of DRG axons to reach unheard of lengths of up to 10 cm *in vitro* ^6-9,11,12^. Building on this technique, we have used stretch-grown axons as the backbone of tissue engineered “living scaffolds”, which to date have been comprised of living sensory axon tracts spanning several centimeters. We have shown that these tissue engineered axonal tracts serve as direct pathways for host axon regeneration by mimicking the developmental action of “pioneer” axons ^6,12,13^. However, it is evident that axons from other neuronal populations, specifically spinal motor neurons, are able to withstand an equal magnitude of mechanical forces as DRG axons during development. However, it is unclear whether this characteristic can be recapitulated under culture conditions for spinal motor neurons, because specific features unique to DRG neurons/axons may endow resiliency under artificial stretch growth conditions, such as their innate robustness and/or the physical architecture of the ganglia.

Therefore, in the current study we developed a facile method of forced cell aggregation, which is used to mimic the architecture of DRG that we predict will increase the tolerance of more fragile neurons and axonal tracts to mechanical forces. Here, engineered micro-spheres of motor neurons were generated from a dissociated cell solution using inverted pyramid wells and simple centrifugation techniques ^14,15^. We have also successfully applied this method of aggregation to skeletal myocytes to create a co-culture system consisting of phenotypically specific populations. By combining these elements, we describe a versatile system in which motor neurons/axons are cultured in a highly controlled manner with either sensory neurons or myocytes and mechanically stretch grown to form long engineered axonal tracts. This system shows promise as an anatomically and physiologically relevant tool to study development, disease, and cell-drug interactions *in vitro*, and can also be applied as reparative constructs to facilitate the reconstruction and/or regeneration of neuro-myo connections following trauma or neurodegeneration *in vivo*.

## Methods

All procedures are approved by the Institutional Animal Care and Use Committees at the University of Pennsylvania and the Michael J. Crescenz Veterans Affairs Medical Center and adhered to the guidelines set forth in the NIH Public Health Service Policy on Humane Care and Use of Laboratory Animals.

### 2.1 Dorsal Root Ganglion Harvest and Culture

Dorsal root ganglia (DRG) were obtained from embryonic day 16 (E16) Sprague Dawley (Charles River) pups. The mother rat was euthanized with CO_2_ asphyxiation followed by decapitation. All subsequent cell harvesting steps until plating were performed on ice or on a cold block. Pups were extracted from the ovaries and immersed in a 10 cm petri dish containing cold L-15 media (Life Technologies). The pups were decapitated, and organs were extracted ventrally. The vertebral column was cut open, and the spinal cords were harvested from the ventral side. The spinal cord was placed in a 35mm petri dish containing cold Hanks Balanced Salt Solution (HBSS) (Life Technologies). Whole DRG explants were plucked from the spinal cord using fine surgical forceps and placed in a 1.5mL conical tube containing 1.2mL L-15 media. DRG were plated on poly-D-lysine (PDL) and laminin coated culture surfaces. Specifically, cell culture surfaces were treated with 20 μg/mL PDL diluted in sterile cell culture sterile water (Lonza) overnight. The next day, culture surfaces were rinsed three times with cell culture grade water to wash away excess PDL and allowed to incubate in 20 μg/mL Laminin for at least 2 hours. The laminin solution was removed from the culture substrate, and DRG were plated in culture dishes flooded with DRG plating media consisting of Neurobasal medium (Life Technologies), supplemented with 2% B-27, 1% fetal bovine serum, 0.5 mM L-Glutamine, 20 ng/mL nerve growth factor, 2.5 g/L glucose, and 40 μM mitotic inhibitors to inhibit glial cell proliferation.

### 2.2 Rat Motor Neuron Harvest and Forced Neuronal Aggregate Culture

Motor neurons were harvested from the spinal cord of E16 Sprague Dawley rat embryos. Culture plates were prepared as described above for rat DRG culture. All harvest procedures prior to dissociation were conducted on ice. After the pups were decapitated and tails were snipped, the vertebral column was cut open from the dorsal surface, and spinal cords were extracted and placed in cold HBSS media. The meninges and any remaining DRG were then removed from the cord.

The spinal cord was placed in 2.5% 10X trypsin diluted in 1mL L-15 for 15 mins at 37°C with intermittent agitation every 5 mins. After dissociating the spinal cord, the trypsin solution was removed, carefully aspirating to avoid disturbing the tissue, and 100 μL of 1mg/mL DNAse and 4% BSA in 900 μL L-15 was added. The suspension was triturated, and the supernatant placed in a sterile 15mL tube, taking care not to disturb the digested tissue. L-15 media was added to the solution to bring the final volume of the extracted supernatant to 10 mL by adding L-15 media. After adding the L-15 media, a 4% BSA cushion was added to the bottom of the tube using a glass pipette. The suspension was centrifuged at 280g for 10 minutes. Additional dissociation media, consisting of 20 μL 1 mg/mL DNase and 50 μL 4% BSA in 900 μL L-15, was added to the original cell solution. Following trituration, the supernatant was placed in a separate sterile tube taking care not to disturb the tissue. This dissociation process was repeated 2-3 times ^16^. The supernatant was aspirated from the first cell dissociation following centrifugation, and the cell pellet was combined with the cell suspension obtained from the repeated DNase and BSA dissociation process. The final volume was brought to 10 mL by adding L-15, and a 1mL layer of Optiprep density gradient was added to the bottom of the tube. The cell suspension was centrifuged for 15 min at 520g and 4°C. Following centrifugation, the cells at the interface between the Optiprep layer and media were collected and suspended in 5mL L-15 with a 500 μL 4% BSA cushion at the bottom. Cells were centrifuged again at 280g for 10 min at 25°C. Following centrifugation, the supernatant was discarded, and cells were resuspended in motor neuron plating media consisting of glial conditioned media. To condition the media, Neurobasal media containing 10% FBS was added to a flask of spinal astrocytes and incubated overnight. The next day, the media was taken out and supplemented with 37ng/mL hydrocortisone, 2.2 μg/mL isobutylmethylxanthine, 10 ng/mL BDNF, 10 ng/mL CNTF, 10 ng/mL CT-1, 10 ng/mL GDNF, 2% B-27, 20ng/mL NGF, 20 μM mitotic inhibitors, 2 mM L-glutamine, 417 ng/mL forskolin, 1 mM sodium pyruvate, 0.1 mM β- mercaptoethanol, 2.5 g/L glucose ^16^. The cells were then plated at a density of approximately 4-5 x 10^4^ cells/cm^2^ on PDL and laminin and coated surfaces, as described for DRGs. To create motor neuron aggregates, dissociated cells were plated in “pyramid” wells ^14,15^. These are wells comprised of polydimethyl siloxane (PDMS) in a hollow, inverted “pyramid” shape, with the cells gathering at the “tip” of the pyramid (Fig. 4B). Cell suspension volume of 12 μL was added to each pyramid, and centrifuged at 1500 RPM for 5 min. The wells were then flooded with motor neuron plating media and incubated for 24 hours to allow the aggregates to form. For fluorescent labeling of the aggregates, cells were incubated overnight in media consisting of AAV-GFP (AAV1.hSynapsin.EGFP.WPRE.bGH, UPenn Vector Core) or AAV-mCherry (AAV1-CB7-CI-mCherry.WPRE.rBG, UPenn Vector Core vector. The following day, the aggregates were extracted from the well using a pipette and plated in the desired culture dish (Fig. 4B).

### 2.3 Myocyte Culture and Aggregation

Mouse skeletal myoblast cell line (C2C12) were cultured in tissue culture flasks in growth media consisting of DMEM-high glucose (Gibco) + 20% FBS + 1% Penstrep). Cells were allowed to reach 90% confluency before being maintained in differentiation media (DMEM-high glucose + 1% normal horse serum + 1% Penstrep) for 5 days to allow differentiation into myocytes and subsequent formation of elongated myofibers. The cells were detached from the tissue culture flasks by trypsin treatment (0.5% Trypsin-EDTA for 15 min). Growth media was added, and the cell suspension was centrifuged at 300g for 5min. The cell pellet was dissolved in growth media such that the cell concentration was approximately 6.5×10^6^ cells/ml. PDMS based pyramid wells as described above were used to form cell aggregates. Specifically, 12µl of cell solution was added to each pyramid well and centrifuged at 1500 RPM for 5mins. The PDMS inserts were filled with growth media comprising of AAV9-tMCK-GFP for transduction and incubated for 24 hours to allow cell aggregation.

### 2.4 Use of Custom-Built Mechanobioreactors

The stretch-growth bioreactors are composed of a custom designed expansion chamber, linear motion table, stepper motor, and controller (Fig. 1). The expansion chamber serves as a tissue culture environment consisting of a sealed enclosure with a port for CO_2_ exchange, removable carriage designed to slowly separate two populations of neuronal somata and connecting rods to allow for displacements (Fig. 1D-K). Attached to the carriage is a bottom substrate (base Aclar) made of optically transparent Aclar 33C film (198 µm thick) that remains stationary and an overlapping movable substrate (towing membrane; ∼10 µm thick) (Fig. 1I-K). The latter is placed on top of the base Aclar. Two populations of cells were plated adjacent to each other, one on the base Aclar and one on the towing membrane and allowed to extend processes and form connections. The towing membrane was then moved in a controlled manner via an automated microstepper motor and controller system with a computer-controlled LabView (National Instruments) user interface (Fig. 1B-D), thus separating the stationary population of cells (base Aclar) from the moving population of cells (towing membrane). The result was two populations of neuronal somata separated by a defined distance and spanned by axon fascicles (Fig. 1J, K).

**Figure 1:**
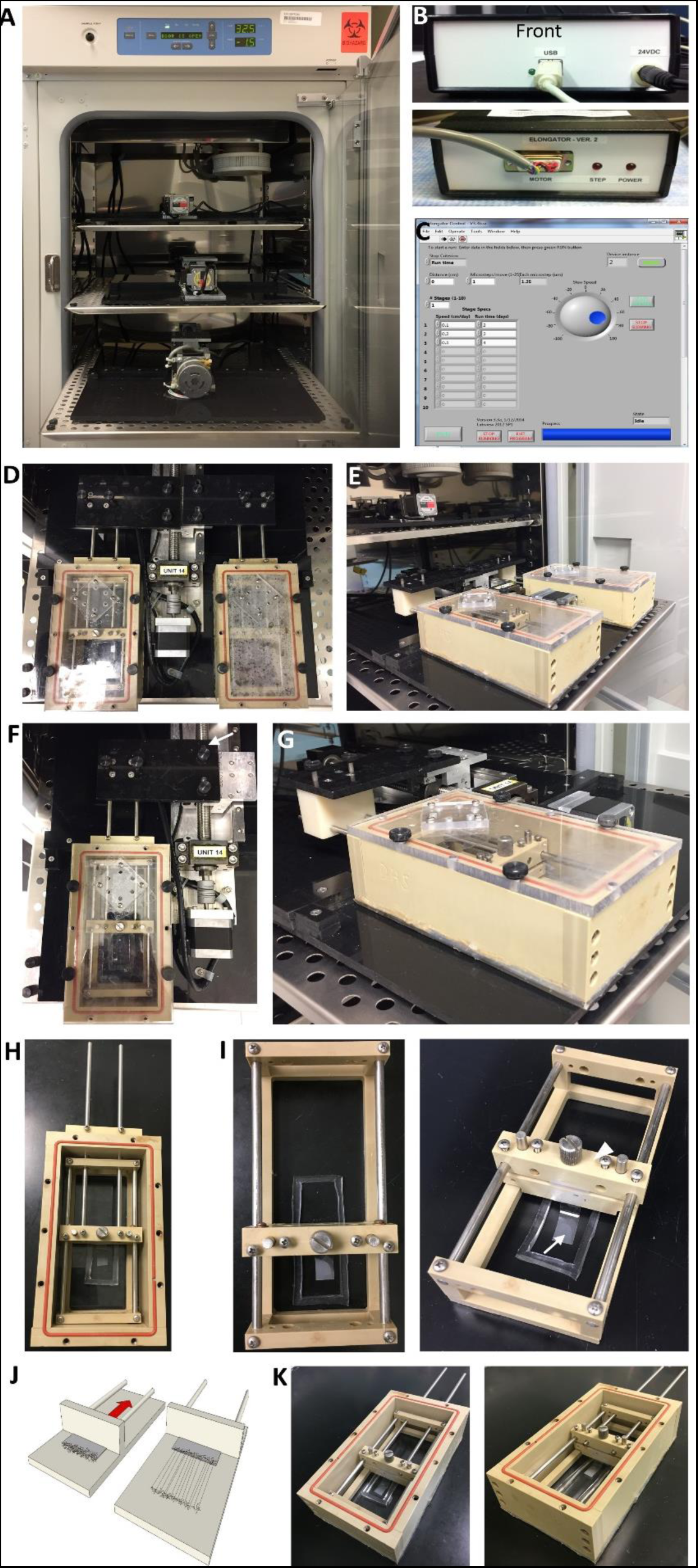
Stretch apparatus set up. (A) Dehumidified incubator with stretch tables containing stepper motors. (B) Control box front and back with connections going from the stretch table to the control box (DIN cable) and from the control box to the computer (USB). (C) Screenshot of the LabView program to control the stretch tables. The speed and duration of mechanical tension may be adjusted. (D) Two custom-built expansion chambers can be attached to one stepper motor. (E) Side view of the attached expansion chambers. (F) Top view of an expansion chamber connected to the stepper motor with an attachment (white arrow). (G) Side view of expansion chamber connected to stepper motor. (H) Carriage consisting of towing membrane and rods to connect to the stepper motor inside of an expansion chamber. (I) Carriage with towing membrane (arrow) adhered to the towing block (arrow head) that will be pulled back by the rods (depicted in H). (J) Schematic representation of cells plated along towing membrane that is connected to the towing block with axon stretch growth when the towing block is pulled back due to continuous mechanical forces. (K) Depiction of towing block pulled back within an expansion chamber.

Elongator Aclar substrates (base and towing membrane) were treated in 1N NaOH for 24 hours to increase hydrophilicity of the substrate, rinsed in deionized water, and then attached to the stretching frame or carriage using medical grade RTV silicone adhesive. The glue-assembled mechanobioreactors were exposed to UV in the hood for 48 hours to allow for sterilization. All culture surfaces were first treated with 20 µg/mL PDL followed by a 20 µg/mL laminin solution, as described above.

### 2.5 Axon Stretch-Growth

#### Dorsal Root Ganglion Sensory Neurons

Mechanobioreactors were prepared as described above, and DRG were manually plated in 2 straight rows approximately 1 mm apart on either side of the interface of the towing membrane and base Aclar using forceps to position DRGs at the desired location. Approximately 12 DRG were placed on each side of a 1 cm wide towing membrane. The DRGs were then allowed to adhere for 3-4 hours on a warming pad heated to 37°C before being moved into the incubator. Axonal networks were allowed to form between the two populations of DRG for 5 days. On day 5, DRGs were transduced with AAV-GFP+ vector. The vector was added to the media for 24 hours, after which time it is washed away through a complete media change prior to initiating stretch. For 1 cm stretch, mechanical tension was applied for 10 days at 1 mm/day or for 2 days followed by 2 mm/day for 4 days. A half media change was completed once every week. Once axons were elongated to the desired length, the culture was removed and stored in a normal humidified incubator until needed.

#### Spinal Cord Motor Neurons

Motor neuron aggregates were plated in custom-built mechano bioreactors prepared as described above for DRGs. Bioreactors were filled with plating media and aggregated neurons were plated in two rows on either side of the towing membrane approximately 0.5 mm apart without touching. Approximately 10 aggregates were plated on each side of a 1 cm towing membrane approximately 500 μm apart and incubated in the bioreactors for 6 days to allow axonal connections to form between the two populations. The bioreactor was then connected to the stepper motor within a dedicated non-humidified incubator (5% CO_2_ at 37°C) using an adapter for application of mechanical tension on the axons (Fig. 1A, D-G). The adapter slid on to the metal rods connected to the towing block and attached directly to the stepper motor (Figure 1D). A half media change was done once per week while neurons were in culture. As with DRG neurons, cultures were stored in a normal humidified incubator once the desired length had been reached.

#### DRG Sensory Neurons + Spinal Cord Motor Neurons

DRG were harvested as described previously and plated in mechanobioreactors along the towing membrane at 50% density to leave space for motor neuron aggregates. AAV-mCherry vector was added to the media and allowed to incubate overnight to produce red fluorescent sensory neurons and axons. Motor neuron aggregate formation was completed the following day, with motor neurons expressing GFP, as described above, to differentiate them from sensory DRG. Media in the mechanobioreactor was replaced with fresh motor neuron media, as was the media in the pyramid wells to rid traces of viral vector. Motor neuron aggregates were added to the mechanobioreactors. Mixed motor sensory TENGs were plated in a way that sensory and motor aggregates were alternating; approximately 5-6 DRG and 5-6 motor neuron aggregates were plated on either side of the towing membrane, resulting in 10-12 ganglia and aggregates total on each side. It should be noted that motor aggregates were plated about 0.5 mm apart from each other across the towing membrane, whereas DRG were plated approximately 1 mm apart from each other. As with pure motor constructs, cultures were allowed to incubate for 6 days to allow axonal connections to form across the towing membrane. After 6 days *in vitro* (DIV), mechanical tension was applied to the cultures at a rate of 1 mm/day.

#### Spinal Cord Motor Neurons + Skeletal Myocytes

Myocyte aggregates were plated on the base Aclar membrane and were cultured in differentiation media for 2 days in the mechanical bioreactors described above. Subsequently, motor neuron aggregates were plated on the towing membrane and the cells were maintained in serum-free motor neuron plating media for 7 days to allow axonal connections to form with the myocytes. Mechanical tension was applied at a rate of 0.5 mm/day to obtain stretch grown axons connected to myocyte aggregates.

### 2.6 Immunocytochemistry and Imaging

Sensory neuron, motor neuron and myocyte cultures were routinely imaged using phase contrast microscopy techniques on a Nikon Eclipse Ti inverted microscope with Nikon Elements Basic Research software. Immunocytochemistry techniques were performed as previously described ^12^. Cultures were fixed in 4% formaldehyde for 30 min, rinsed in phosphate buffered saline (PBS), and permeabilized using 0.3% Triton X100 plus 4% horse serum for 60 min. Primary antibodies used to identify sensory DRG and motor neurons were added (in PBS + 4% serum solution) at 4°C for 12 hrs. Mouse anti-β-tubulin III (Sigma C8198) was used to identify a specific microtubule protein expressed in neurons, sheep anti-choline acetyltransferase (ChAT; abcam ab18736) and rabbit anti-p-75 (Sigma N3908) were used as specific motor neuron markers, and rabbit anti-calcitonin gene related peptide (CGRP; Sigma C8198) was used as a marker for sensory neurons and axons. After rinsing, secondary antibodies (1:500 in PBS + 4% NHS) were applied at room temperature for 2 hours (Alexa 561 donkey anti-rabbit IgG and Alexa 488 donkey anti-mouse IgG). Stretched or non-stretched motor and/or sensory cultures were fluorescently imaged using a laser scanning confocal microscope (Nikon A1 Confocal Microscope). For each culture, multiple confocal z-stacks were digitally captured and analyzed.

### 2.7 Group Sizes, Data Quantification, and Statistical Analyses

Conditions were optimized for motor neuron culture, stretch growth, and co-culture with sensory neurons or skeletal myocyes. Health and phenotype of cells was qualitatively assessed, while axon density and fascicle width were quantified. In order to determine the effect of cell density on motor neuron aggregate diameter, the diameter of aggregates comprising of 12.5×10^3^, 25×10^3^, 50×10^3^, 75×10^3^, 100×10^3^ and 120×10^3^ cells (n = 8 each) was measured using FIJI software.

Once stretch grown, the axon/fascicle width and area of axon coverage was measured for pure sensory (n=10), pure motor (n=18), and mixed motor-sensory (n=13) constructs. Phase contrast images of stretch grown constructs were acquired, and the diameter of all discernible axons and fascicles were measured using FIJI software. A grid was overlayed on the image and the diameter of all the axons and fascicles was taken equidistant from the edge of the cell body region. Additionally, a histogram was created to show the frequency distribution of axon/fascicle width in sensory, motor, and mixed motor-sensory constructs. To quantify the percentage of axon coverage over a given area (1 cm), using the same grid overlay, the length of areas within one construct lacking axon or fascicle outgrowth equidistant from the edge of the cell body region were measured and summed. The following percent difference equation was used:

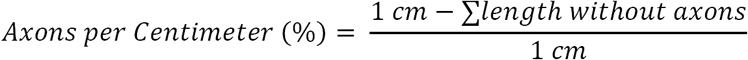

One way ANOVA followed by Tukey’s multiple comparison test (Graphpad Prism) was used to test statistical significance (p < 0.5).

## Results

### 3.1 Development and Characterization of Spinal Motor Neuron Cultures and Spinal Motor Neuron – Sensory DRG Co-Cultures

Once dissociated spinal motor neurons were plated, they would naturally begin to form small clusters at approximately 2 DIV, and by 4 DIV “node-like” structures of adjacent cell body clusters connected by axon tracts had formed (Fig. 2A). These persisted throughout culture out to at least 21 DIV (Fig. 2B). Motor neurons were also co-cultured with whole DRG explants, exhibiting extensive neurite outgrowth and network formation. Additionally, stark differences in size between DRG explants and motor neuron “nodes” self-formed from individual motor neuron somata were observed, with both neuronal subtypes projecting healthy axons (Fig. 2C, D). Neuronal phenotype was confirmed through positive expression of β-tubulin-III (Fig. 2B-D).

**Figure 2:**
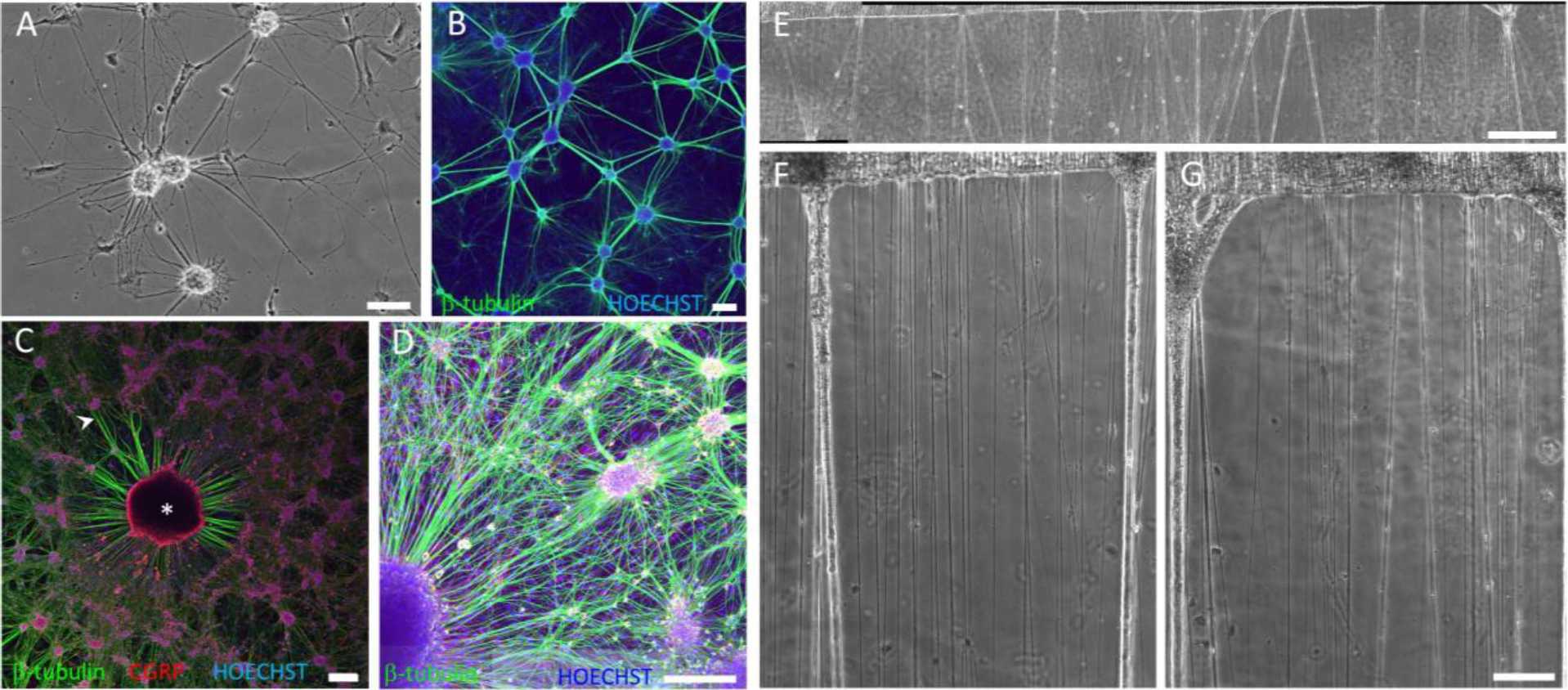
Stretch growth of dissociated motor neurons produces robust motor axons, however at a slow rate of displacement. (A) Phase contrast image of dissociated motor neurons plated on PDL-Laminin coated polystyrene at 4 days *in vitro* (DIV). Scale: 250 μm. (B) Confocal reconstruction of dissociated motor neuron cultures at 21 DIV expressing β-tubulin (green) and the nuclear counterstain HOECHST (blue). Note the self-formation of small aggregated networks of motor neurons at both time points. Scale: 250 μm. (C) Confocal reconstruction of whole DRG explant (*) surrounded by dissociated motor neuron co-culture positively labeled for β-tubulin (green) and the sensory neuron marker, CGRP (red), and HOECHST nuclear counterstain (blue). Arrow head denotes sensory DRG axons. Scale: 250 μm. (D) DRG whole explant (*) surrounded by dissociated motor neuron co-culture clearly showing the size differential between motor neuron self-formed nodes and whole DRG (β-tubulin-III, green; HOECHST, blue). Scale: 250 μm. (E) Phase contrast image of stretch grown dissociated spinal motor neurons stretched to 1 cm at a rate of 0.1 mm/day. Scale: 1000 μm. (F, G) Higher magnification of stretch grown motor axons, showing robust motor axons. Scale: 100 μm.

### 3.2 Demonstration of Axonal Stretch-Growth from Spinal Motor Neurons

After establishment of successful spinal motor neuron cultures, the next step was “stretch-growth” of motor axon tracts. Since optimal stretch parameters for DRG neurons have previously been determined and used to generate stretch-grown sensory axons spannng several centimeters^6-9^, we adapted and modified these parameters for use in motor neuron culture. First, dissociated motor neurons were plated and mechancial tension was applied to the culture using our custom-built mechanobioreactors. These motor axons demonstrated healthy stretch-growth, but only at extremely slow rates of displacement (0.1-0.3 mm/day) (Fig. 2E-G). When faster rates of displacement were attempted, leading to higher strain rates, the axons could not respond to the stress and snapped (Fig. 3B, F-H). In contrast, DRG are able to tolerate much higher strain rates, as they are able to withstand initial displacement rates of greater than 1 mm/day (Fig. 3A, C-E). Although we demonstrated that motor axons projecting from dissociated neurons could be stretch-grown, the slow stretch growth rate necessary to sustain axon continuity in these dissocated motor neurons was undesirable as the required time frame was both inefficient and may be too long to maintain neuronal health (approximately 1 cm axon tracts in 1-3 months). However, it was not apparent if the slower rate of motor axon stretch-growth was due to an inherent growth limitation of motor axons or if the 3-D, aggregated nature of DRG explants conferred an advantge (e.g., structural and/or physiological) for stretch-growth.

**Figure 3:**
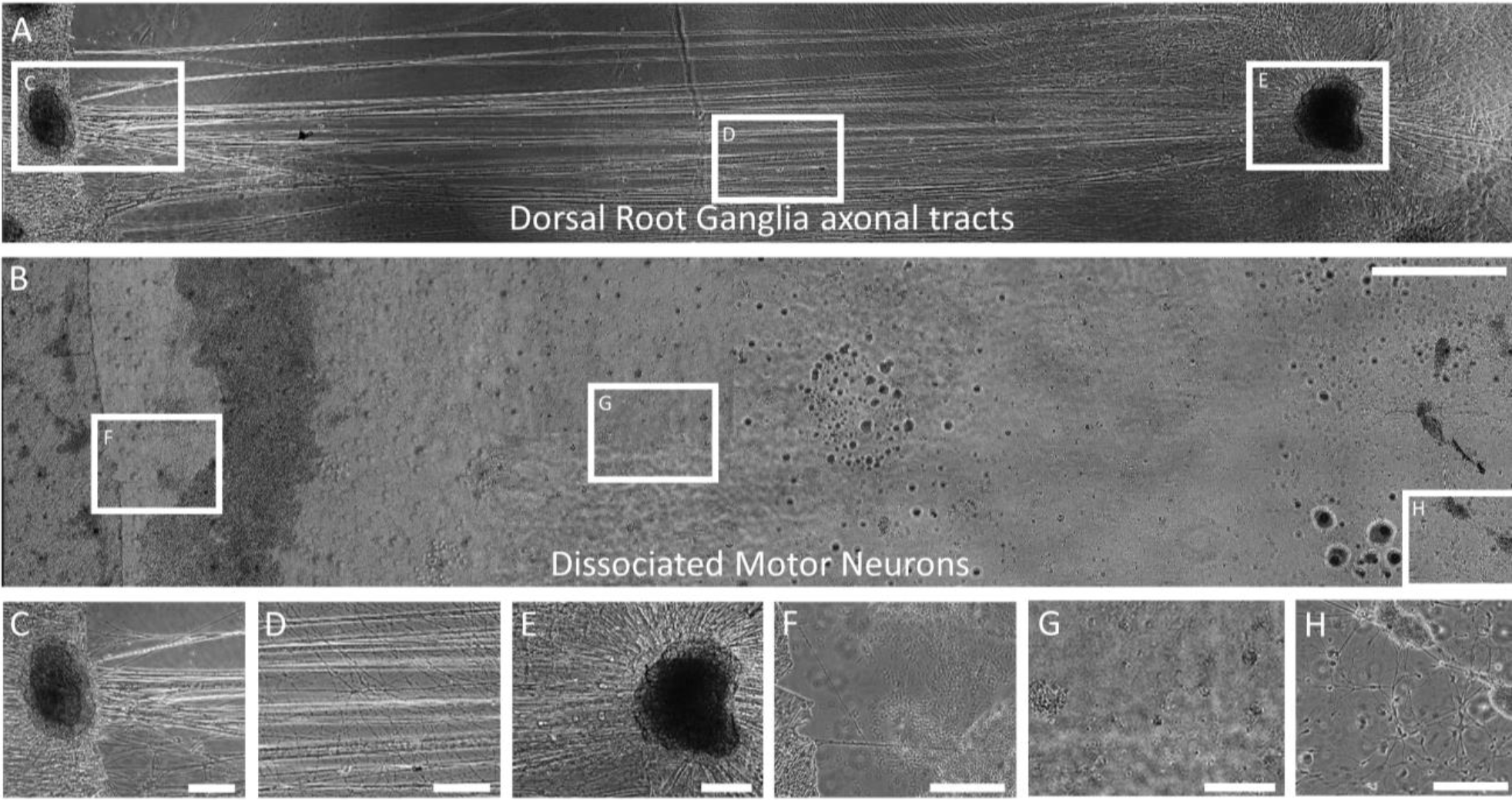
Axons from dissociated motor neurons were unable to withstand a rate of displacement that was tolerated by axons from DRG explants. Phase contrast images showing that application of mechanical tension (continuous displacement of 1.0 mm/day) to axonal networks from (A) whole DRG explants produced robust axonal tracts spanning 1 cm. (B) When an equal rate of displacement was applied to dissociated motor neurons, the axonal networks were unable to withstand the force and snapped. Scale: 1000 μm. Higher magnification showing the contrast between robust axons from (C-E) DRG explant stretch growth and (F-H) the lack of stretch grown axons when dissociated spinal motor neurons were subjected to an equivalent displacement rate as whole DRG. Scale: 250 μm.

### 3.3 Forced Aggregation of Spinal Motor Neurons

To address this issue, our goal was to develop a method that woud mimic the architecture of DRG explants using spinal motor neurons. Here, motor neurons were acquired, dissociated, and purified using density gradients as described above. Then, we created “forced neuronal aggregates” by implementing a facile method utilizing PDMS inverted pyramid wells and gentle centrifugation (Fig. 4A, B, D). When motor neurons were aggregated, long, robust process outgrowth was seen surrounding the entire cell body core (Fig. 4E,F), whereas non-aggregated, dissociated neurons exhxibited much smaller neurite outgrowth (Fig. 4C). The diameter of aggregated neuronal cultures can be controlled by changing the density of cells within the aggregates. As expected, the density of cells in the aggregate was directly proportional to the diamater of the aggregate (Fig. 4G). However, it should be noted that with too large of a diameter, the aggregates were more susceptible to disassembling during the plating process and/or developing a necrotic core. Aggregation did not effect cell phenotype as the cells were confirmed to be motor neurons by labeling for motor neuron markers, such as ChAT and the nerve growth factor receptor p75, along with the general neuronal marker β-tubulin III (Fig. 4H-O). Forced aggregation of motor neurons resulted in healthy clusters of cells exhibiting robust axon outgrowth, with controlled variability in aggregate diameter.

**Figure 4:**
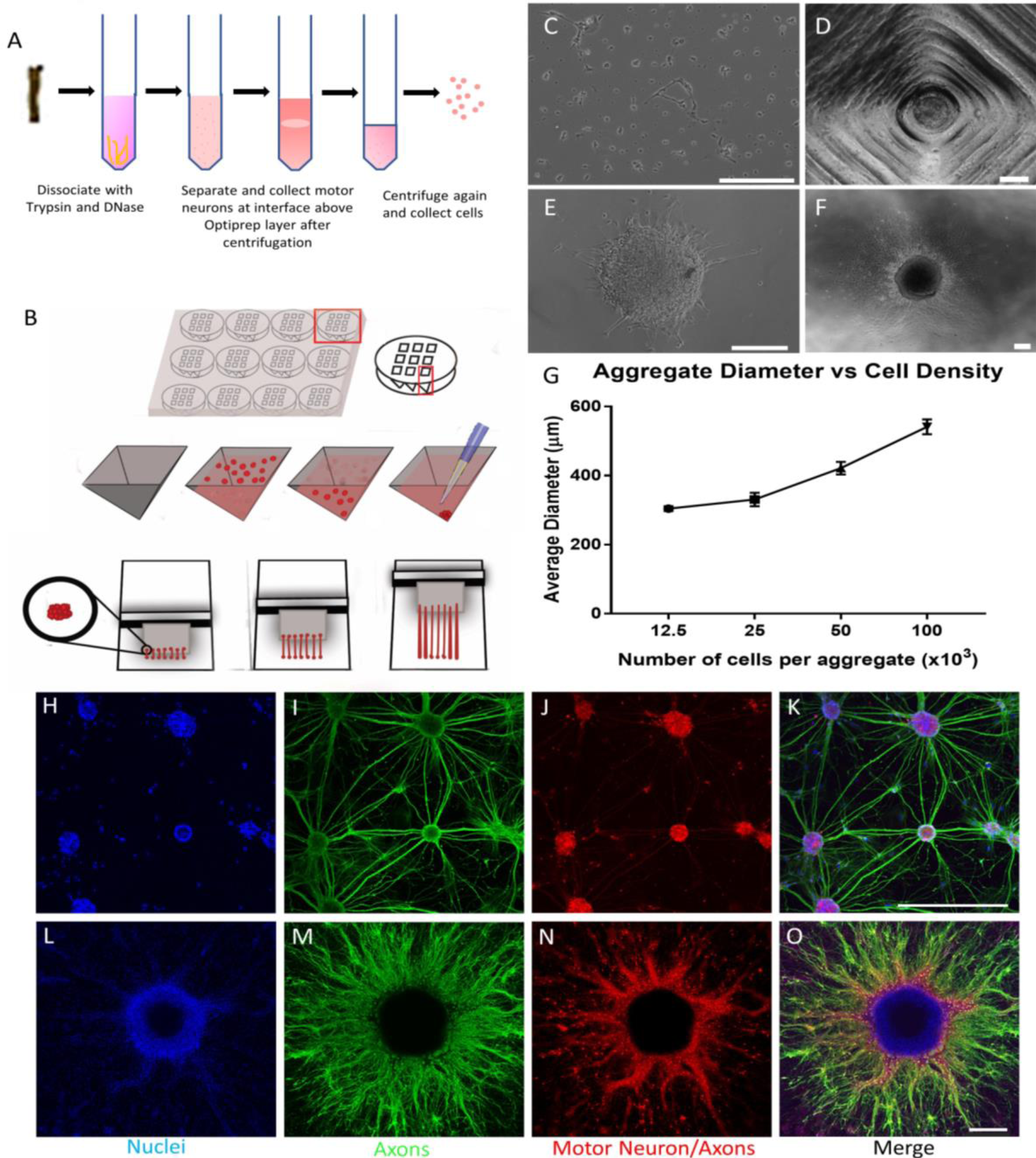
Forced neuronal aggregation and stretch growth methodology. Motor neuron dissociation, culture, and stretch growth. (A) Motor neurons were harvested from embryonic rat spinal cords and dissociated using bovine serum albumin (BSA) and Optiprep density gradients to acquire a more pure population of motor neurons and (B) plated in custom-built pyramid shaped wells, centrifuged, and incubated in plating media overnight to allow aggregates of motor neurons to form. (C) Dissociated motor neurons exhibited short and more sparse neurites at 1 DIV. (D) Motor neuron aggregate 24 hours after centrifugation in an inverted pyramid well. Aggregated motor neurons at (E) 1 DIV and (F) 5 DIV exhibiting longer and more robust neurite outgrowth. (G) The diameter of the aggregate was proportional to the cell density of the aggregate. Both (H-K) dissociated and (L-O) aggregated motor neurons positively labeled for (H,L) nuclear marker (HOECHST, blue); (I,M) neuronal marker (β**-**tubulin-III, green) and (J,N) motor neuron-specific marker (ChAT, red) at 7 DIV, with (K,O) showing all channels merged. (C-F) scale bars: 250 μm; (H-O) Scale bars: 500 μm

### 3.4 Optimization of Motor Axon Stretch-Growth Using Motor Neuron Aggregates

Our goal was to apply mechanical tension at rates equivalent to those tolerated by DRG axons to force-aggregated motor neuron axons in order to produce intact axon tracts spanning at least 1 cm. Motor aggregates were plated along two sides of the towing membrane-base interface, as described in the Methods. We found that this neuronal aggregation culture system allowed more robust neurite outgrowth, and thus was able to withstand higher rates of and greater magnitued of displacement. The motor axon network was now able to tolerate displacement rates as high as 1 mm/day, and have been routinely stretched to 1 cm (Fig. 5A,B). Confirmation of motor neuron/axon phenotype was also seen (Fig. 5C-J). Of note, motor axons have been stretched to 1.7 cm to date, with greater lengths expected (Fig. 5K-O).

**Figure 5:**
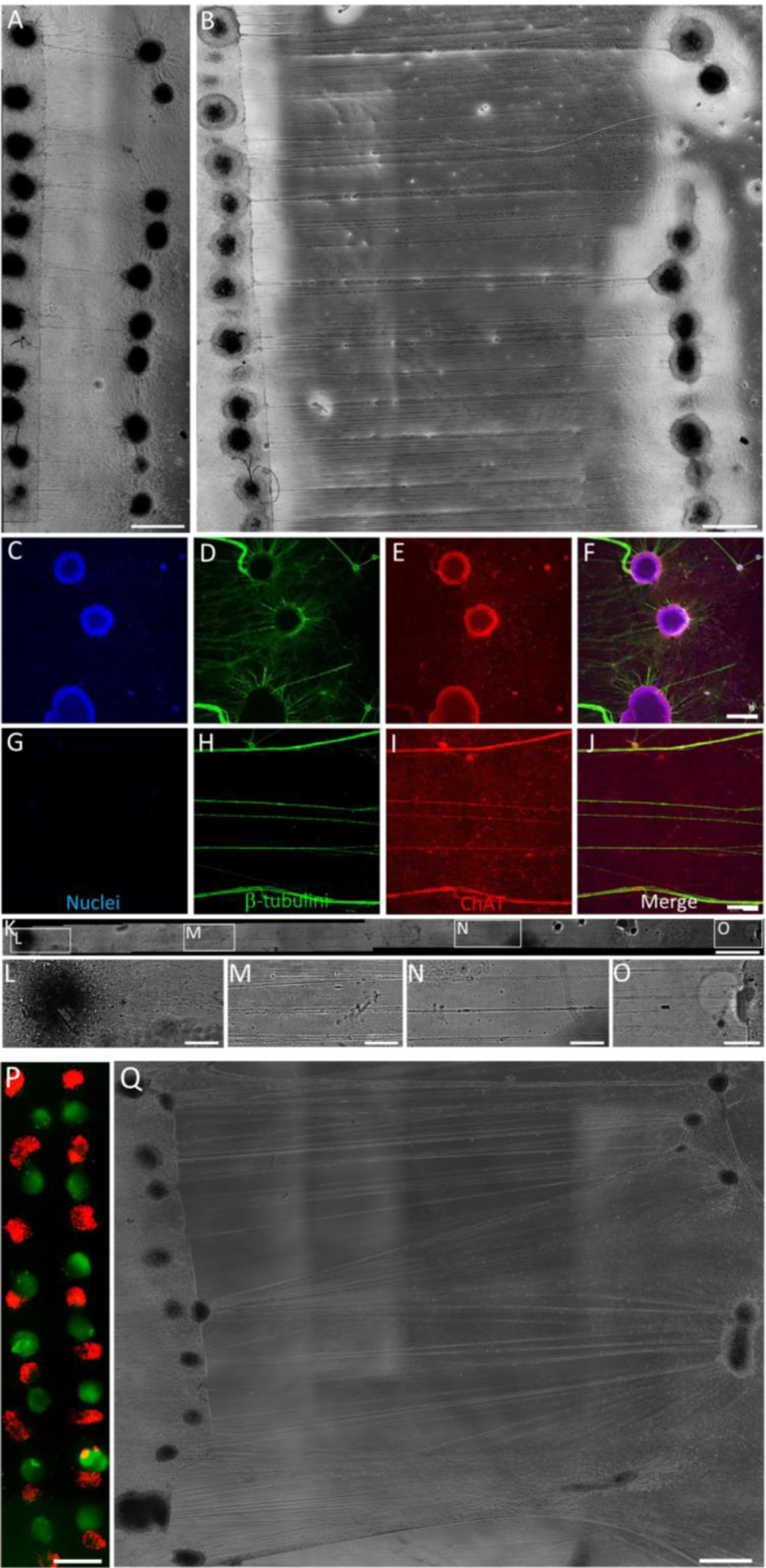
Development and characterization of stretch-grown motor axon constructs. Rat motor neurons were isolated from spinal cords and forced into neuronal aggregates. The neuronal aggregates were plated in custom-built mechanobioreactors, and tension was applied to the axons at a rate of 1 mm/day. (A) Phase contrast image after tension was applied for 1 day, and axons have stretched to approximately 1 mm. Scale: 1000 μm. (B) Phase contrast image after the motor axons have stretched to 1 cm. Scale: 1000 μm. Confocal reconstruction of motor neuron forced aggregate (C-F) cellular region and (G-J) pure axonal section, showing (C,G) nuclear stain (HOECHST, blue), (D,H) axons (β**-**tubulin-III, green), (E, I) motor neuron specific marker (p75, red), and (F, J) merge of all channels. Scale: 500 μm. (K) Motor axons stretched to 1.7 cm. Scale: 1000 μm. (L-O) Phase contrast zoom-in images show healthy stretch grown motor axons across the entire 1.7 cm distance. Scale: 250 μm. A mixed motor-sensory construct developed by alternating separately acquired sensory DRG (mCherry-positive, red) and motor neuron aggregates (GFP-positive, green). (P) Fluorescent image prior to application of mechanical tension. (Q) Phase contrast image depicting 1 cm long sensory and motor axons spanning sensory and motor cell body regions. Scale: 1000 μm.

In addition to optimizing stretch parameters, the effect of mixed sensory and motor neuron/axon cultures was of interest since pure sensory TENGs have shown promise upon transplantation in PNI models *in vivo*, but axon regeneration is believed to be modality dependent. These mixed motor-sensory neuronal cultures consisted of alternating DRG explants and aggregated motor neurons, keeping the ratio of DRG to motor neuron aggregates equal (Fig. 5P). The aggregated motor neurons and DRG were differentially labeled prior to co-culture in the mechanobioreactor (Fig. 5P). Even in the co-culture system, robust neurite outgrowth of both sensory and motor axons was observed, leading to stretch growth of dense, fascicularized axons out to at least 1 cm, with much longer axon tracks possible (Fig. 5Q). Overall, the novel method of forced neuronal aggregation resulted in motor neuron structures that were resilient to higher rates of displacement, resulting in long, robust stretch grown motor axons. Additionally, the motor neuron aggregates could be co-cultured with sensory DRG explants to produce mixed motor-sensory constructs consisting of stretch grown sensory and motor axons.

### 3.5 Comparison of Motor, Sensory, and Mixed Motor-Sensory Axon Stretch-Growth

Once pure motor and mixed motor-sensory stretch-grown constructs were generated, our objective was to compare axon/fascicle health and morphology between construct-types. Motor neuron only, sensory neuron only, and mixed motor-sensory neuron cultures were generated and subjected to axonal stretch-growth under identical conditions. Overall neuron and axon health was similar across the three types of constructs (Fig. 6A-C). Following axon stretch-growth to 1 cm, we found that motor axon constructs exhibited a statistically lower density of axon fascicles than sensory only or mixed motor-sensory constructs (Fig. 6G). This was likely due to the fact that stretched motor aggregates produce an increased number of significantly thicker fascicles than sensory only or mixed motor-sensory stretched axons (Fig. 6H). As expected, mixed constructs consisted of axons with attributes similar to both sensory only and motor only constructs. Namely, there was an increased number of wider axon fascicles as was similar to motor only constructs, as well as thinner axons resembling sensory only constructs (Fig. 6D-F).

**Figure 6:**
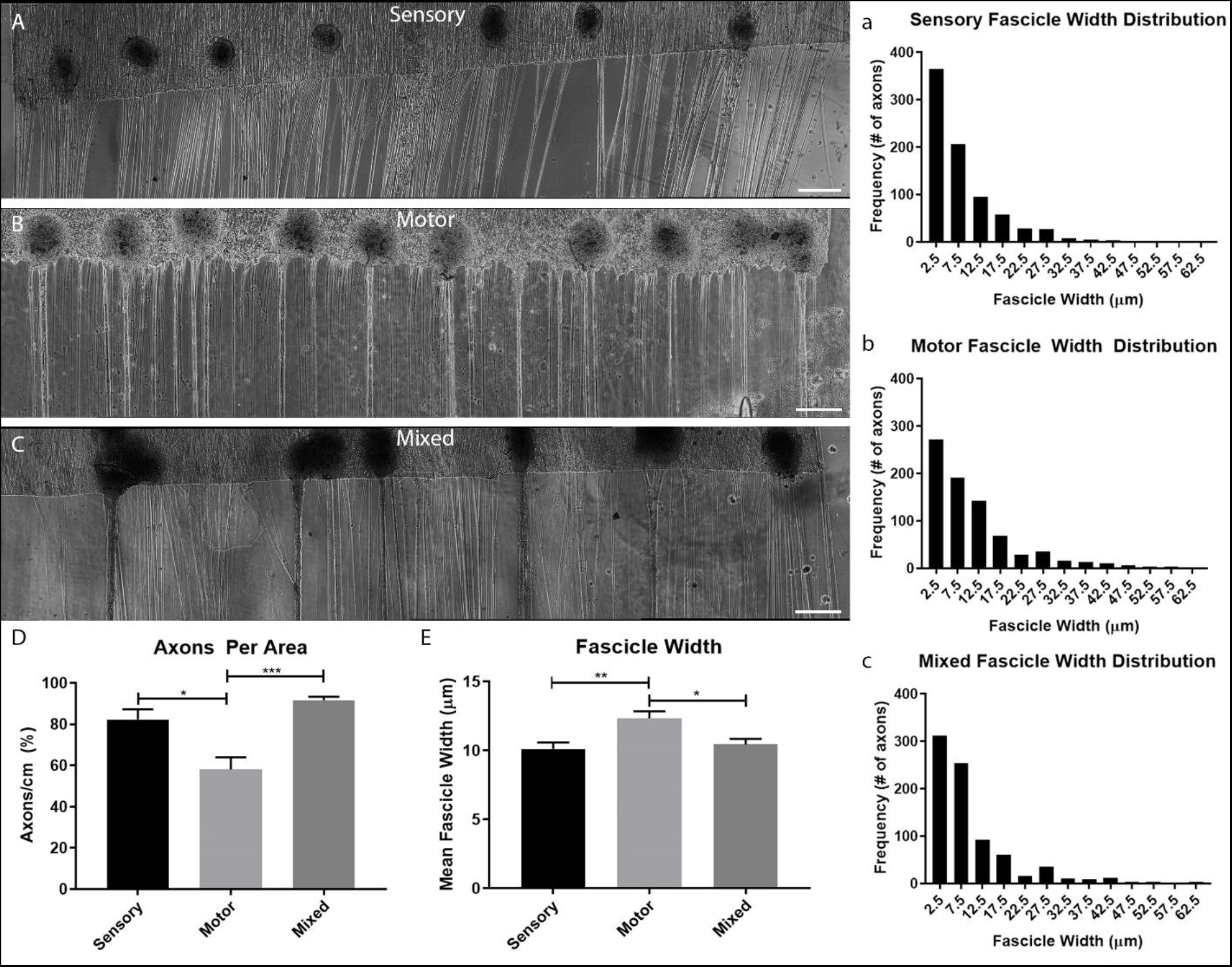
Differences in motor, sensory, and mixed modality constructs. Three types of constructs comprising 1 cm axon growth from (A) sensory neurons only, (B) motor neurons only, and (C) mixed sensory + motor neurons, were generated by implementing culture of neuronal aggregates in custom-built mechanobioreactors. Histograms depict differences in fascicle width between (a) sensory, (b) motor, and (c) mixed axonal constructs. (D) Axon density was significantly higher in mixed and sensory constructs as compared to motor only constructs (*p<0.05, ***p<0.001). (E) Likewise, fascicle width was significantly greater in motor in constructs as compared to sensory or mixed constructs. (n = 9 cultures each group; *p<0.05,

### 3.6 Myocyte-Motor Neuron Co-culture and Stretch Growth

Since motor axons innervate muscle *in vivo*, our goal was to develop a system in which motor axons have formed connections with myocytes, but the cell bodies of each remain separate, thus better mimicking physiological conditions. Multi-nucleated aligned myofibers were formed after 12 DIV in differentiation media (Fig. 7A-D). Aggregates formed from predifferentiated myocytes behaved as phenotypically specific 3-D spheroids projecting myofibers (Fig. 7E-G). Co-culture of myocyte aggregates with spinal motor neuron aggregates resulted in long axons projecting from the motor neuron aggregate to innervate neighboring myofibers (Fig. 7H-J). Within the mechanobioreactors, myocyte and motor neuron aggregates were observed to form connections after approximately 7 DIV. The motor neuron aggregates growing on the towing membrane were then gradually displaced, producing long axons that were observed to be projecting from the motor neuron aggregates and connected to the myocytes (Fig. 7K).

**Figure 7:**
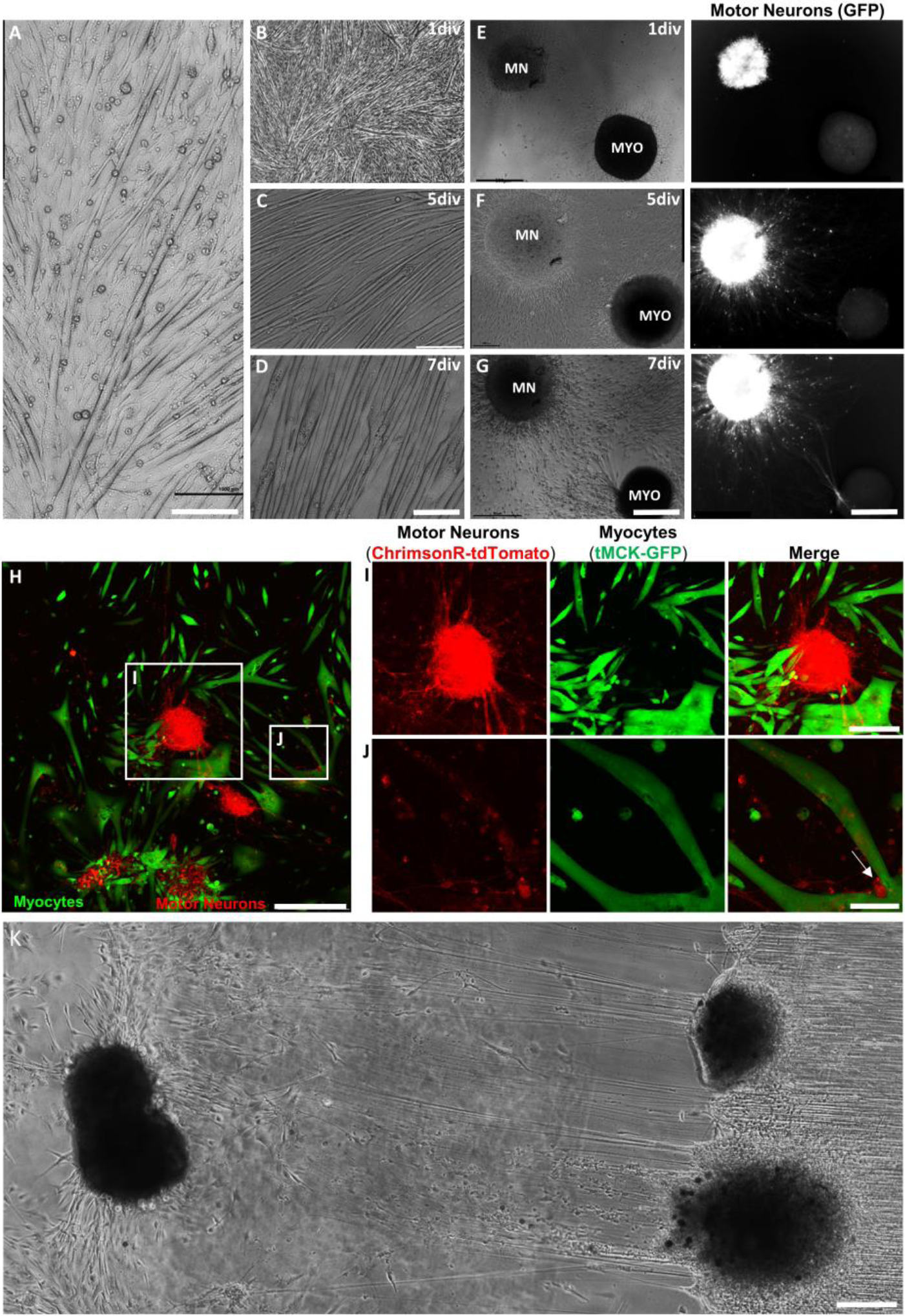
Culture and forced aggregation of myocytes produced robust aligned myofibers *in vitro*. (A) Mouse skeletal myoblast cell line (C2C12) differentiated to form aligned, elongated myofibers by 12 DIV. Scale: 1000 μm. (B-D) The skeletal myocytes progressively fused with each other and align to form multinucleated myofibers. Scale: 500 μm. (E-G) Forced aggregates of pre-differentiated myocytes were co-cultured with spinal motor neuron aggregates (transduced with AAV1-hSyn-GFP) and axonal outgrowth towards mycoytes was observed. Scale: 500 μm. (H) Confocal microscopy of motor neuron-myocyte aggregated co-culture showed spinal motor neuron aggregates transduced with AAV1-hSyn-ChrimsonR-tdTomato (red) sending out long axons to innervate myofibers expressing muscle creatine kinase (green). Scale: 500 μm. (I,J) Zoom-in images show motor neuron-myocyte interactions, with the arrow indicating points of innervation in the myofibers. Scale: 250 μm. (K) Phase contrast images clearly showing axons spanning 3 mm from motor aggregates (*) to myocyte aggregates (arrowhead). Scale: 250 μm

## Discussion

Injury and disorders of the spinal cord and peripheral nervous system are increasingly common and can lead to significant or complete loss of sensory and/or motor function. Neuronal cell bodies are housed in the spinal column and project nerves comprising of bundles of axons to the rest of the body. Sensory DRG as well as motor neurons are located in or around the spinal cord and project long axons that innervate the periphery. Due to the long distances these axons span, it has proved extremely difficult to recapitulate lost nerve or create a suitable “bridge” to promote regeneration of axon tracts following large nerve defects spanning several centimeters. Repair using the current gold standard, the autologous nerve graft (autograft) requires taking healthy sensory nerve to repair damaged nerve, and yields about 50% recovery in smaller defects, but remains largely ineffective for major peripheral nerve injury (i.e. spanning ≥5 cm). Additionally, SCI is common and may lead to severe injury of axon tracts and subsequent disruption of signal transmission. Approximately 50% of SCIs are diagnosed as complete SCIs, affecting both sides of the body equally, and often leading to complete loss of function due to loss of spinal motor neurons as well as ascending and descending axonal tracts.

To address these challenges, repair strategies that are able to replace and/or repair lost neurons and/or axonal pathways are necessary. Here, neural tissue engineering techniques may be useful, especially those utilizing 3D cell culture to better mimic the *in vivo* conditions such as the microenvironmental and cellular interactions underlying growth, maturation, and function of various cell types. Numerous techniques for 3D culture have been developed, including culturing cells on a biomaterial or biologically-derived scaffold, culturing cells within a liquid matrix followed by polymerization of the supporting material, or with only cells, negating the use of a scaffold material ^17,18^. Incorporation of a scaffold to provide structure and support for cells to grow is widely used. However, 3D culturing of cells without the use of support material is deemed more challenging. This technique is commonly utilized to create tumor models, where a hanging drop culture is used to create aggregates of cells to mimic the tumor microenvironment ^19^. It has also recently been used to create aggregates of stem cells that can be used to study development or produce uniform aggregates of stem cells that differentiate into cardiomyocytes ^14,20^ and to create spheroids for paracrine factor secretion ^15^. Many of these spheroids are developed in suspended culture to pre-activate cells prior to contact with host cells to increase secretion of desirable growth factors ^21^, to model embryonic development ^22^, and to study differentiation into neurons and glia ^23^. In contrast to these approaches, we developed a method of forced neuronal aggregation where aggregates are attached to the substratum *in vitro* during formation. As described above, our method of forced aggregation allowed us to engineer motor neuron “ganglia” that were more robust when subjected to mechanical forces capable of inducing stretch-growth.

Applying this technique, we extended our previous findings of axon stretch-growth of pure sensory axons to include pure motor and mixed motor-sensory stretch grown axons. Here, we developed novel constructs exhibiting regions of aggregated neuronal soma spanned by long, aligned axonal tracts. We found that axons spanning two populations of aggregated motor neurons can tolerate mechanical tension and have been stretched to approximately 2 cm, with longer axon lengths likely attainable. Additionally, the co-culture of motor neurons and DRG neurons produced robust neurite outgrowth and fasciculation of stretch-grown axons of both neuron types, with motor-sensory fascicles appearing significantly thicker than pure sensory fascicles, and comparable to the thickness of pure motor fascicles. These data suggest that TENGs can be comprised solely of sensory neurons (DRG), motor neurons (MN), or mixed-modality (DRG+MN) neurons, thus potentially providing a tailored approach for targeted, modality-specific nerve repair.

In our experiments, several parameters had to be optimized to achieve long aligned motor axonal tracts, similar to the sensory tracts seen for DRG. These parameters include the time that neurons were in culture as well distance between motor aggregates before application of mechanical tension, rate at which external mechanical tension was applied, the length to which the axons were stretched, and co-culture with sensory neurons and myocytes. In addition, the cell concentration, aggregate size and inter-aggregate distance were some of the crucial parameters that effected stretch growth. We found that aggregated motor neuron cultures with approximatelly 4-5×10^4^ cells in each aggregate yielded aggregates of diameter similar to that of DRG. However, it should be noted that most aggregates plated for stretch consisted of 8-10×10^4^ cells as they exhibited more robust axon outgrowth. These motor neuron aggregates were plated at an optimal distance of 0.5 mm apart from each other which allowed axons to form robust connections with the aggregate(s) on the other side of the towing membrane interface. The distance that motor neuron aggregates were plated was approximately half of the distance that DRGs were plated from each other. This is due to the slower rate at which axons extend from motor neuron somata and the less dense initial axonal projections compared to DRG axons. However, some distance was necessary as motor aggregates were seen to fuse to the corresponding aggregate on the other side of the towing membrane interface when plated too closely together. As seen previously, an increased rate of displacemnet may be applied to axons over time as the axons lengthened and became acclimated to the external application of mechanical tension ^24^. Table 1 compares stretch rates tolerated by different neuronal cell types with varying architecture *in vitro*. Interestingly, other groups have developed internal fixator devices that are able to lengthen nerves after injury *in vivo* by applying strains of approximately 20% to the proximal side of the damaged nerve before reattaching to the distal nerve stump ^25^.

**Table 1:** Rate of displacement tolerated by rat neural cells in vitro.

| Neural Source & Structure |  | Rate of Displacement (mm/day) |  |  |
| --- | --- | --- | --- | --- |
| Subtype | Architecture | Day 1-2 | Day 3-4 | Day 5+ |
| Dorsal Root Ganglia | Whole Ganglia | 1.0 | 2.0-3.0 | 4.0<sup>8</sup> |
| Motor Neuron | Dissociated | 0.1 | 0.3 | 0.5 |
| Motor Neuron | Aggregate | 1.0 | 1.0<sup>#</sup> | 1.0<sup>#</sup> |
| Motor Neuron – Dorsal Root Ganglia | Aggregate – Ganglia | 1.0 | 1.0<sup>#</sup> | 1.0<sup>#</sup> |
| <sup>#</sup>higher rates were not attempted in the current study |  |  |  |  |

In our *in vitro* studies, we found that dissociated motor neurons were only able to tolerate a rate of displacement that is approximately 10% of that which DRG and aggregated motor neuron axons were able to withstand. This may be due to the increased fasciculation of motor axons in aggregated form as compared to dissociated cultures, allowing for increased resistance to external force. The increase in fasciculation was very clearly seen with thicker axonal tracts in aggregated stretched motor constructs when compared to dissociated stretched motor constructs. Interestingly, motor constructs exhibited thicker fascicles but the lowest density of axon fascicles (Fig. 6B). This may be due to conservation of mass; since the fascicles were thicker, the axon density appeared lower and may be mistaken as less healthy. However, in reality, these constructs were likely equally healthy or healthier than sensory constructs due to the thicker, more robust fascicles. In contrast, sensory constructs exhibit many more individual axons or smaller bundles of axons, but increased axon density within a given area due to the axons growing in a more dispersed fashion (Fig. 6A). Mixed constructs exhibit characeristics of both sensory and motor constructs. Some of the fascicles, such as the ones stemming from motor aggregates, were thicker and appeared more sparse - as seen in the histogram with some axon fascicles present with larger diameters; whereas bundles of axons sprouting from DRG were much thinner and continuously cover a larger area - as seen with the distribution of axon fascicles with smaller axons (Fig. 6C).

This system produces tracts of motor axons that are able to better recapitulate nerves *in vivo* by generating axons that are longer than any other known *in vitro* technique. This is the first demonstration of generating an *in vitro* system in which stretch-grown motor axon tracts connect separate populations of motor neurons and muscle cells. Due to their morphological resemblance to host nerve, these axon tracts can be used as an *in vitro* testbed to further study three key areas of neural tissue engineering - development, homeostasis, and disease. The axon tracts can be used to determine mechanisms of stretch growth and axon-facilitated axon regeneration, with a focus on cell-cell interactions during development; neuron/axon function and regulation of homeostasis; and neurodegenerative diseases of the PNS such as amyotrophic lateral sclerosis (ALS) and spinal muscular atrophies, among others. A more holistic model consisting of phenotypically separated populations of myocytes and these motor neurons with long, aligned axons can be used as a model to study innervation and muscle functions or for implant following injury, as the system, consisting of a cluster of motor neurons with long axons growing into muscle cells, better preserves anatomical fidelity. For regenerative purposes, these aggregated motor neuron constructs can be implemented as living scaffolds comprised of motor and/or sensory tissue engineered nerve grafts for directed bridging following peripheral nerve trauma. In conjuction with PNI, these constructs can be tansplanted into the spinal cord following injury to replace lost motor neurons with long axonal extensions spanning a meter or more to distal targets.

## 4. Conclusion

This is the first demonstration of mechanical elongation of spinal cord motor axons that may have applications as anatomically- and physiologically-relevant testbeds for neurophysiological, developmental, and/or pathophysiological studies. This technology could prove invaluable in furthering our understanding of the mechanisms of nerve injury and subsequent regeneration to better develop therapies that promote more effective recovery. Additionally, we have built on previous reports of axonal stretch-growth by applying this technique in conjunction with engineered micro-aggregates of muscle cells to create biologically inspired tissue engineered constructs that recapitulates the architecture of long projecting motor axons integrated with mature skeletal myocytes. Tissue engineered motor axon or motor axon-myocyte complexes can serve as an *in vivo*-like model to further study long distance axonal conduction, neuromuscular junction formation and function, as well as the role of innervation in muscle tissue maturation. Collectively, these axonal constructs can also be used as a clinical repair or replacement strategy by acting as tissue engineered “living scaffolds” to exploit axon-facilitated axon regeneration to drive long distance, modality specific axonal regeneration following PNI or SCI. In addition, the engineered nerve-muscle complexes may also be applied in regenerative medicine by creating pre-innervated muscle coupled to long-projecting spinal motor axons to restore neuro-myo connections via direct replacement and integration following neuromuscular injury or trauma.

## Acknowledgements

Financial support was provided by the U.S. Army Medical Research and Materiel Command [W81XWH-15-1-0466 (Cullen) & W81XWH-16-1-0796 (Cullen)], the National Institutes of Health [U01-NS094340 (Cullen) & F31-NS090746 (Katiyar)], the National Science Foundation [Graduate Research Fellowships DGE-1321851 (Struzyna)], and the Department of Veterans Affairs [BLR&D Merit Review I01-BX003748 (Cullen)].

## References

1 Fabricius, C., Berthold, C. H. & Rydmark, M. Dimensions of individual alpha and gamma motor fibres in the ventral funiculus of the cat spinal cord. J Anat 184 (Pt 2), 319–333 (1994).

2 Sepp, K. J., Schulte, J. & Auld, V. J. Peripheral glia direct axon guidance across the CNS/PNS transition zone. Developmental biology 238, 47–63 (2001).

3 Cullen, D. K., Vukasinovic, J., Glezer, A. & Laplaca, M. C. Microfluidic engineered high cell density three-dimensional neural cultures. J Neural Eng 4, 159–172, doi:10.1088/1741-2560/4/2/015 (2007).

4 Cullen, D. K., Wolf, J. A., Smith, D. H. & Pfister, B. J. Neural tissue engineering for neuroregeneration and biohybridized interface microsystems in vivo (Part 2). Crit Rev Biomed Eng 39, 241–259 (2011).

5 Irons, H. R. et al. Three-dimensional neural constructs: a novel platform for neurophysiological investigation. J Neural Eng 5, 333–341, doi:10.1088/1741-2560/5/3/006 (2008).

6 Huang, J. H. et al. Long-term survival and integration of transplanted engineered nervous tissue constructs promotes peripheral nerve regeneration. Tissue engineering. Part A 15, 1677–1685, doi:10.1089/ten.tea.2008.0294 (2009).

7 Loverde, J. R., Tolentino, R. E. & Pfister, B. J. Axon stretch growth: the mechanotransduction of neuronal growth. Journal of visualized experiments: JoVE, doi:10.3791/2753 (2011).

8 Pfister, B. J., Iwata, A., Meaney, D. F. & Smith, D. H. Extreme stretch growth of integrated axons. J Neurosci 24, 7978–7983, doi:10.1523/JNEUROSCI.1974-04.200424/36/7978 [pii] (2004).

9 Smith, D. H., Wolf, J. A. & Meaney, D. F. A new strategy to produce sustained growth of central nervous system axons: continuous mechanical tension. Tissue engineering 7, 131–139, doi:10.1089/107632701300062714 (2001).

10 Huang, J. H. et al. Harvested human neurons engineered as live nervous tissue constructs: implications for transplantation. Journal of neurosurgery 108, 343–347 (2008).

11 Higgins, S., Lee, J. S., Ha, L. & Lim, J. Y. Inducing neurite outgrowth by mechanical cell stretch. Biores Open Access 2, 212–216, doi:10.1089/biores.2013.0008 (2013).

12 Katiyar, K. S., Winter, C. C., Struzyna, L. A., Harris, J. P. & Cullen, D. K. Mechanical elongation of astrocyte processes to create living scaffolds for nervous system regeneration. J Tissue Eng Regen Med 11, 2737–2751, doi:10.1002/term.2168 (2017).

13 Struzyna, L. A., Katiyar, K. & Cullen, D. K. Living scaffolds for neuroregeneration. Curr Opin Solid State Mater Sci 18, 308–318, doi:10.1016/j.cossms.2014.07.004 (2014).

14 Ungrin, M. D., Joshi, C., Nica, A., Bauwens, C. & Zandstra, P. W. Reproducible, ultra high-throughput formation of multicellular organization from single cell suspension-derived human embryonic stem cell aggregates. PLoS One 3, e1565, doi:10.1371/journal.pone.0001565 (2008).

15 Zimmermann, J. A. & McDevitt, T. C. Pre-conditioning mesenc*hymal stromal cell spheroids for immunomodulatory paracrine factor secretion. Cytotherapy 16, 331–345, doi:10.1016/j.jcyt.2013.09.004 (2014).

16 Graber, D. J. & Harris, B. T. Purification and culture of spinal motor neurons from rat embryos. Cold Spring Harb Protoc 2013, 319–326, doi:10.1101/pdb.prot074161 (2013).

17 Edmondson, R., Broglie, J. J., Adcock, A. F. & Yang, L. Three-dimensional cell culture systems and their applications in drug discovery and cell-based biosensors. Assay Drug Dev Technol 12, 207–218, |p|doi:10.1089/adt.2014.573 (2014).

18 Haycock, J. W. 3D cell culture: a review of current approaches and techniques. Methods Mol Biol 695, 1–15, doi:10.1007/978-1-60761-984-0_1 (2011).

19 Timmins, N. E. & Nielsen, L. K. Generation of multicellular tumor spheroids by the hanging-drop method. Methods Mol Med 140, 141–151 (2007).

20 Bauwens, C. L., Toms, D. & Ungrin, M. Aggregate Size Optimization in Microwells for Suspension-based Cardiac Differentiation of Human Pluripotent Stem Cells. Journal of visualized experiments: JoVE, doi:10.3791/54308 (2016).

21 Bartosh, T. J. et al. Aggregation of human mesenchymal stromal cells (MSCs) into 3D spheroids enhances their antiinflammatory properties. Proc Natl Acad Sci U S A 107, 13724–13729, doi:10.1073/pnas.1008117107 (2010).

22 Hookway, T. A., Butts, J. C., Lee, E., Tang, H. & McDevitt, T. C. Aggregate formation and suspension culture of human pluripotent stem cells and differentiated progeny. Methods 101, 11–20, doi:10.1016/j.ymeth.2015.11.027 (2016).

23 Imrik, P. & Madarasz, E. Importance of cell-aggregation during induction of neural differentiation in PCC-7 embryonal carcinoma cells. Acta Physiol Hung 78, 345–358 (1991).

24 Loverde, J. R. & Pfister, B. J. Developmental axon stretch stimulates neuron growth while maintaining normal electrical activity, intracellular calcium flux, and somatic morphology. Front Cell Neurosci 9, 308, doi:10.3389/fncel.2015.00308 (2015).

25 Vaz, K. M., Brown, J. M. & Shah, S. B. Peripheral nerve lengthening as a regenerative strategy. Neural regeneration research 9, 1498 (2014).

